# Coordination of di-acetylated histone ligands by the ATAD2 bromodomain

**DOI:** 10.1101/2021.07.23.453464

**Authors:** Chiara M. Evans, Margaret Phillips, Kiera L. Malone, Marco Tonelli, Gabriel Cornilescu, Claudia Cornilescu, Simon J. Holton, Mátyás Gorjánácz, Liping Wang, Samuel Carlson, Jamie C. Gay, Jay C. Nix, Borries Demeler, John L. Markley, Karen C. Glass

## Abstract

The ATPase Family, AAA domain-containing protein 2 (ATAD2) bromodomain (BRD) has a canonical bromodomain structure consisting of four α-helices. ATAD2 functions as a co-activator of the androgen and estrogen receptors as well as the MYC and E2F transcription factors. ATAD2 also functions during DNA replication, recognizing newly synthesized histones. In addition, ATAD2 is shown to be up regulated in multiple forms of cancer including breast, lung, gastric, endometrial, renal, and prostate. Furthermore, up-regulation of ATAD2 is strongly correlated with poor prognosis in many types of cancer, making the ATAD2 bromodomain an innovative target for cancer therapeutics. In this study, we describe the recognition of histone acetyllysine modifications by the ATAD2 bromodomain. Residue-specific information on the complex formed between the histone tail and the ATAD2 bromodomain, obtained through nuclear magnetic resonance spectroscopy (NMR) and X-ray crystallography, illustrates key residues lining the binding pocket, which are involved in coordination of di-acetylated histone tails. Analytical ultracentrifugation, NMR relaxation data, and isothermal titration calorimetry further confirm the monomeric state of the functionally active ATAD2 bromodomain in complex with di-acetylated histone ligands. Overall, we describe histone tail recognition by ATAD2 BRD and illustrate that one acetyllysine group is primarily engaged by the conserved asparagine (N1064), the “RVF” shelf residues, and the flexible ZA loop. Coordination of a second acetyllysine group also occurs within the same binding pocket, but is essentially governed by unique hydrophobic and electrostatic interactions making the di-acetyllysine histone coordination more specific than previously presumed.

## Introduction

ATPase family, AAA domain-containing protein 2 (ATAD2), also known as ANCCA (AAA+ nuclear coregulator cancer◻associated), has a bromodomain and an AAA ATPase domain [1]. The ATAD2 bromodomain (BRD) contains the canonical, left-handed bundle structure of four α-helices (αZ, αA, αB, αC), which support the two highly variable loops, ZA (residues 1003-1030) and BC (residues 1063-1068), which are responsible for forming peptide docking sites [2,3]. The bromodomain of the ATAD2 protein is classified as a family IV bromodomain based on conserved residues and structural features it shares with other members of family IV [2]. This chromatin binding protein is able to recognize acetylated lysine through its bromodomain, a chromatin reader domain that is highly conserved across species [4,5]. Bromodomains are known to bind ε-N acetylated lysines [6]. The ATAD2 BRD is known to recognize the N-terminal tail of histone H4 that is acetylated on lysine 5 (H4K5ac) and lysine 12 (H4K12ac), as well as histone H3 at lysine 14 (H3K14ac), and the structures of the ATAD2 BRD in complex with these ligands are available in the Protein Data Bank (PDB) (PDB IDs: 4TT2, 4QUT, and 4TT4) [3,7]. The structure of the ATAD2 BRD with H3K14ac is unique in that the ligand is bound on the outside of the bromodomain, making molecular contacts with R1005 and H1076 [7]. The structures of the ATAD2 BRD with H4K5ac and H4K12ac reveal several molecular contacts responsible for coordination and selectivity of acetyllysine ligands. These include a conserved asparagine at position 1064, a conserved tyrosine at position 1021, the gatekeeper residue, isoleucine 1074, and members of the RVF shelf (R1007, V1008, and F1009, known as the WPF shelf in other bromodomains) [3,7]. The function of ATAD2 remains generally unknown, but it has been shown to interact with estrogen the receptor, the androgen receptor, E2Fs, and the MYC oncogene [1,8–10] In addition to these roles, ATAD2 is known known to be up-regulated in multiple different types of cancer, including breast, lung, gastric, endometrial, renal, prostate, and more recently thyroid [9–16]. Up-regulation of ATAD2 is often correlated with poor patient outcomes, and because of this it can be used as prognostic marker [10,15,17,18]. The up-regulation of ATAD2 in a variety of diverse cancer types suggests that pharmacological inhibition of its bromodomain function could be beneficial in cancer therapies, and is currently under investigation by both pharmaceutical companies and research institutes alike [19–21]. More recently, Bayer AG published the structure of BAY-850, which selectively binds to the ATAD2 BRD isoform over the ATAD2B BRD, by using a unique mechanism of action in which it creates dimers of the ATAD2 BRD [22]. Compound 18a from the Northern Institute for Cancer Research also has strong selectivity for the ATAD2 BRD over the bromodomain-containing protein 4 (BRD4) bromodomain [23]. Although the ATAD2 BRD has historically been called “difficult to drug”, a handful of compounds now exist that have sub-100 nM affinity for the ATAD2 BRD and greater than 100-fold selectivity over other bromodomains, including other members of family IV and the conserved ATAD2 paralogue, ATAD2B [20,22–25].

ATAD2 has been shown to recognize a di-acetylated histone H4K5acK12ac modification that is associated with newly synthesized histones. This interaction appears to facilitate the loading of proliferating cell nuclear antigen (PCNA), a cofactor of DNA polymerase, during DNA replication [26]. A more recent study confirmed the interaction of ATAD2 with several mono- and di-acetylated histone ligands, including H4K5acK12ac [27]. While bromodomains are generally known to recognize single acetyllysine modifications, there are several instances in which bromodomains are observed binding to di-acetyllysine. Bromodomains across the 8 families will recognize di-acetylated ligands typically either as a monomer or as a dimer. For example, in family II, the bromodomain testes-associated protein (BRDT) recognizes H4K5acK8ac as a monomer, with both of the acetylation modifications on the histone ligand fitting within the bromodomain’s binding pocket (PDB ID: 2WP2) [28]. In certain instances, the recruitment of the bromodomain is dependent upon the ligand. For example, when BRD4 is bound to H4K5acK8ac, H4K12acK16ac, or H4K16acK20ac, a monomeric interaction is observed (PDB IDs, 3UVW, 3UVX, and 3UVY, respectively) [2]. However, with H4K8acK12ac, a dimer of the BRD4 bromodomain forms to coordinate this ligand (PDB ID: 3UW9) [2]. The bromodomain-containing protein 2 (BRD2) is a different case in that its second bromodomain recognizes H4K5acK12ac via the formation of two homodimers in the asymmetric unit [29]. In family V, the structure of bromodomain adjacent to zinc-finger domain protein 2B (BAZ2B) with ligand H4K8acK12ac is dimeric, with each BRD monomer bound to an acetylation mark simultaneously (PDB ID: 4QC3) [30]. When BAZ2A is bound to H4K16acK20ac, a hydrophobic pocket of residues formed through BRD dimerization coordinates the K20 acetylation (PDB ID: 4QBM) [30].

For the family IV bromodomains, however, only four of the seven members have been shown to bind di-acetylated histone ligands. Bromodomain-PHD finger protein 3 (BRPF3), a paralog of fellow family members BRPF1 and BRPF2/BRD1, has been found to bind a di-acetylated ligand, but the affinity and oligomerization state of this interaction were not published [31]. Interestingly, the structure of the bromodomain-containing protein 9 (BRD9) was solved in complex with the H4K5acK8ac ligand (PDB ID: 4YYI) and appears to form a dimer to coordinate ligand recognition [31]. The gatekeeper residue of BRD9 is a tyrosine, a large aromatic residue that does not allow the binding pocket to accommodate two acetyllysines at once, unlike the gatekeeper residue of BRD4, which a smaller and aliphatic isoleucine residue. BRPF1B was also recently shown to bind the histone H4K5acK8ac ligand as a dimer (PDB ID: 5FFW), where one acetyllysine binds within the bromodomain binding pocket, while the other is flipped upwards towards the second bromodomain at the interface of the two proteins [32]. The interactions between ATAD2 and H4K5acK12ac are necessary for ATAD2’s recruitment to replication sites [27], but there is no structural information available to demonstrate how it coordinates two acetyllysine marks. A molecular dynamics study indicated that coordination of either the K5ac or the K12ac group of the H4K5acK12ac ligand within the ATAD2 BRD binding pocket is almost identical [2]. This also suggests that the ZA and BC loops remain partially disordered even after ligand binding, and this ‘fuzzy’ interaction may promote cooperativity in recognition of the first and second acetyllysines on the histone tail [33].

In this study, isothermal titration calorimetry (ITC), nuclear magnetic resonance (NMR) spectroscopy, analytical ultracentrifugation (AUC), and X-ray crystallography were employed to study and characterize the binding of both mono- and di-acetylated histone ligands to the ATAD2 BRD. Site-directed mutagenesis, coupled with ITC and circular dichroism (CD), were used to further elucidate the key residues involved in di-acetyllysine recognition by the ATAD2 BRD. We found that the ATAD2 BRD recognizes multiple acetylation modifications in a monomeric state, and identified several new hydrogen-bond and hydrophobic interactions that underline the molecular mechanism driving di-acetyllysine recognition.

## Results

### The ATAD2 BRD recognizes multiple acetyllysine marks on histone tails

We recently identified eleven post-translationally modified histone ligands recognized by the ATAD2 bromodomain using the dCypher array technology developed by EpiCypher [27]. To increase our understanding of di-acetyllysine recognition, we used ITC to examine the preferred ligands of the ATAD2 BRD. We characterized the binding affinity of the ATAD2 BRD with various combinations of mono- and di-acetylated peptides from histones H3, H4, and H2A (**Table 1**, **Figure 1**). Our results indicate that the ATAD2 BRD recognizes mono- and di-acetylated histones H4 and H2A, but does not bind to acetylated histone H3 (**Table 1**). Furthermore, the di-acetyllysine histone H4 tail peptides displayed the highest binding affinities. The histone H4K5acK12ac (1-15) ligand bound the strongest (K_D_ = 28.4 ± 1.3 μM), followed by H4K5acK8ac (1-10) (K_D_ = 33.2 ± 5.9 μM), and H4K12acK20ac (1-24) (K_D_ = 41.0 ± 0.7 μM). Histone H4 and H2A ligands with mono-acetylation at lysine 5 are recognized with similar affinities, while mono-acetylation at K8ac is the least favored, with a weak binding affinity (K_D_ = 1130.3 ±121.6 μM). ITC experiments carried out with the ATAD2 BRD mutants (N1064A, I1074A, and I1074Y), demonstrated a complete loss of acetylated histone binding, thus confirming that the conserved asparagine and the “gatekeeper” residues are essential for recognition of both mono-acetylated and di-acetylated histone H4 ligands (**Supplementary Table S1, Supplementary Figure S2**). Circular dichroism (CD) was used to analyze the secondary structure of all the mutants to confirm that the loss of histone tail recognition was not a result of protein misfolding (**Supplementary Table S2**). The unmodified histones H3 and H4, which lack acetylation marks, also displayed no binding (**Table 1**, **Supplementary Figure S1)**.

**Table 1:** Binding affinities and stoichiometry of binding (N) obtained via isothermal titration calorimetry (ITC) for acetylated histone peptide binding to the ATAD2 bromodomain (BRD).

| Histone peptide | Sequence | ATAD2 BRD $K_D$ (μM) | N-value |
| --- | --- | --- | --- |
| H2AK5ac (res 1-12)* | SGRGKacQGGKARA | $38.9 \pm 5.9$ | 0.997 |
| H4 unmodified (res 1-15)* | SGRGKGGKGLGKGGA | No Binding | - |
| H4 unmodified (res 4-17)* | GKGGKGLGKGGA KR | No Binding | - |
| H4K5ac (res 1-10)* | SGRGKacGGKGL | $44.5 \pm 3.3$ | 1.070 |
| H4K5ac (res 1-15)* | SGRGKacGGKGLGKGGA | $39.4 \pm 6.0$ | 1.040 |
| H4K5ac (res 4-17)* | GKacGGKGLGKGGA KR | No Binding | - |
| H4K8ac (res 1-10)* | SGRGKGGKacGL | $1130.3 \pm 121.6$ | 1.144 |
| H4K8ac (res 4-17)* | GKGGKacGLGKGGA KR | No Binding | - |
| H4K12ac (res 1-15) | SGRGKGGKGLGKacGGA | $250.7 \pm 28.3$ | 1.02 |
| H4K12ac (res 4-17)* | GKGGKGLGKacGGA KR | $95.1 \pm 13.1$ | 1.170 |
| H4K16ac (res 10-20) | LGKGGA KacRHRK | No Binding | - |
| H4K5acK8ac (res 1-10)* | SGRGKacGGKacGL | $33.2 \pm 5.9$ | 0.938 |
| H4K5acK12ac (res 1-15)* | SGRGKacGGKGLGKacGGA | $28.4 \pm 1.3$ | 0.910 |
| H4K8acK12ac (res 6-15) | GGKacGLGKacGGA | $392.1 \pm 19.5$ | 1.169 |
| H4K12acK16ac (res 10-20) | LGKacGGA KacRHRK | $302.6 \pm 18.7$ | 1.082 |
| H4K12acK20ac (res 1-24) | SGRGKGGKGLGKacGGA KRHRKacVLRD | $41.0 \pm 0.7$ | 1.006 |
| H3 unmodified (res 1-12)* | ARTKQTARKSTG | No Binding | - |
| H3K9ac (res 1-12) | ARTKQTARKacSTG | No Binding | - |
| H3K14ac (res 8-19)* | RKSTGGKacAPRKQ | No Binding | - |
| H3K18ac (res 8-19) | RKSTGGKAPRKacQ | No Binding | - |
| *Indicates peptide tested in NMR chemical shift perturbation experiments |  |  |  |

**Figure 1:**
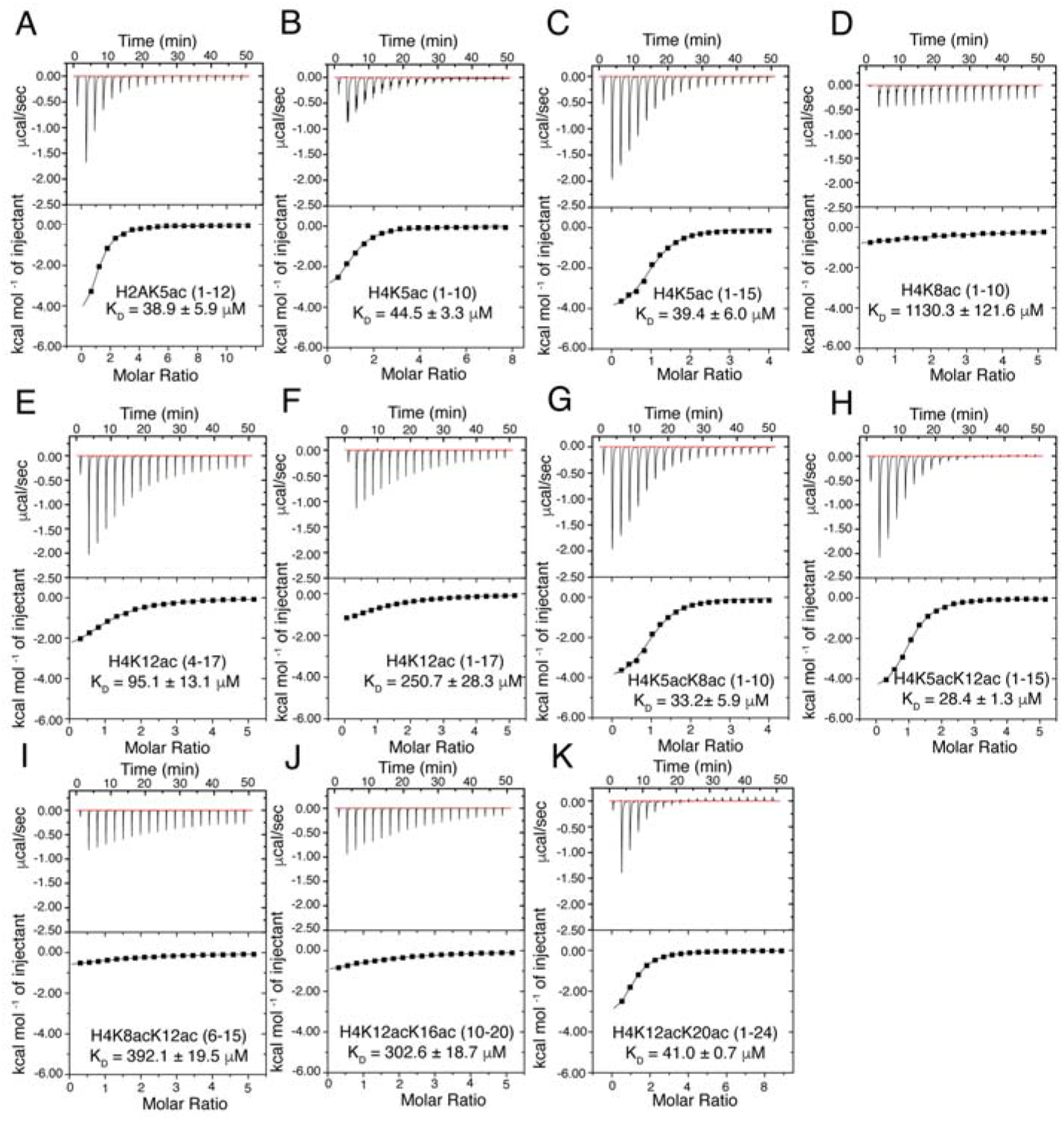
Isothermal titration calorimetry (ITC) measurements for the ATAD2 bromodomain interaction with the histone tail ligands. (A-K) Exothermic ITC enthalpy plots for binding of ATAD2 bromodomain with mono-, and di-acetylated histone ligands. Calculated binding constants and peptides are indicated for each trace.

Recognition of the N-terminal ‘SGR’ residues on the histone H4 tail is essential for coordination with the ATAD2 BRD when acetylation occurs at position 5 or 8, but these residues do not appear to impact coordination of acetyllysine when it is positioned further downstream. This is supported by our ITC data displaying a complete lack of binding to acetylated H4 tail peptides missing the first three residues, e.g., H4K5ac (4-17) and H5K8ac (4-17) (**Table 1**, **Supplementary Figure S1**). However, histone H4K12ac (4-17), binds to the ATAD2 BRD with higher affinity (K_D_ = 95.1 ± 13.1 μM) than the H4K12ac (1-15) peptide (K_D_ = 250.7 ± 28.3 μM). Overall, our ITC data confirm the ATAD2 bromodomain preferentially recognizes di-acetylated histone ligands, and further show that acetyllysine modifications at positions 5 and 12 on the histone H4 tail appear to drive the binding interaction.

### Coordination of di-acetyllysine histone tail by the ATAD2 bromodomain

Selection of the preferred di-acetylated histone H4K5acK12ac ligand by the ATAD2 BRD plays an important functional role in the recruitment of the ATAD2 protein to newly synthesized histones following DNA replications [26]. To further characterize the molecular mechanism used in the selection of histone ligands, we turned to NMR spectroscopy. Chemical shift perturbations (CSPs) induced in the ATAD2 BRD backbone amide resonances upon titrating in mono- and di-acetylated histone ligands were used to delineate the di-acetyllysine histone binding sites in the ATAD2 BRD binding pocket (**Supplementary Figure S3**). Analysis of the NMR titration data established acetylated histone H4 recognition by the ATAD2 BRD with a subset of backbone amide resonances exhibiting significant and consistent CSPs (**Figure 2**). Residues V1008 and F1009 in the RVF shelf, residue K1026 in the ZA loop, residues S1032 and S1033 in the αA helix, the universally conserved N1064, and residues D1068 and G1070 in the BC loop, typically demonstrated large CSPs with most of the histone tail ligands (**Figure 2A, B**). Although the backbone resonances of the majority of the ZA loop residues of the ATAD2 BRD remained unassigned, those of the acetyllysine binding pocket were assigned and could be used to map the CSPs to specific sites. Furthermore, visualization of the acetyllysine histone binding pocket was achieved by mapping the ATAD2 BRD residues displaying highest CSPs onto the apo crystal structure of this bromodomain (PDB ID: 3DAI) (**Figure 3**). Addition of the mono-acetylated histone H4K5ac (1-10) and H4K5ac (1-15) peptides resulted in similar CSP patterns of the ATAD2 BRD residues, with highest perturbations observed for residues in the “RVF” shelf, the ZA loop, and the highly conserved N1064 (**Figure 2A, B**). These indicate that coordination of the K5ac group is preserved and independent of the C-terminus of the histone peptide. The NMR data correlate well with the ITC binding analysis for ATAD2 N1064A mutant, which showed no binding interaction with mono- or di-acetylated histones (**Supplementary Table S1, Supplementary Figure S2**). Similarly, mutation of the “gatekeeper” residue I1074 to either alanine or a bulkier tyrosine, also resulted in complete loss of acetyllysine recognition (**Supplementary Table S1, Supplementary Figure S2**).

**Figure 2:**
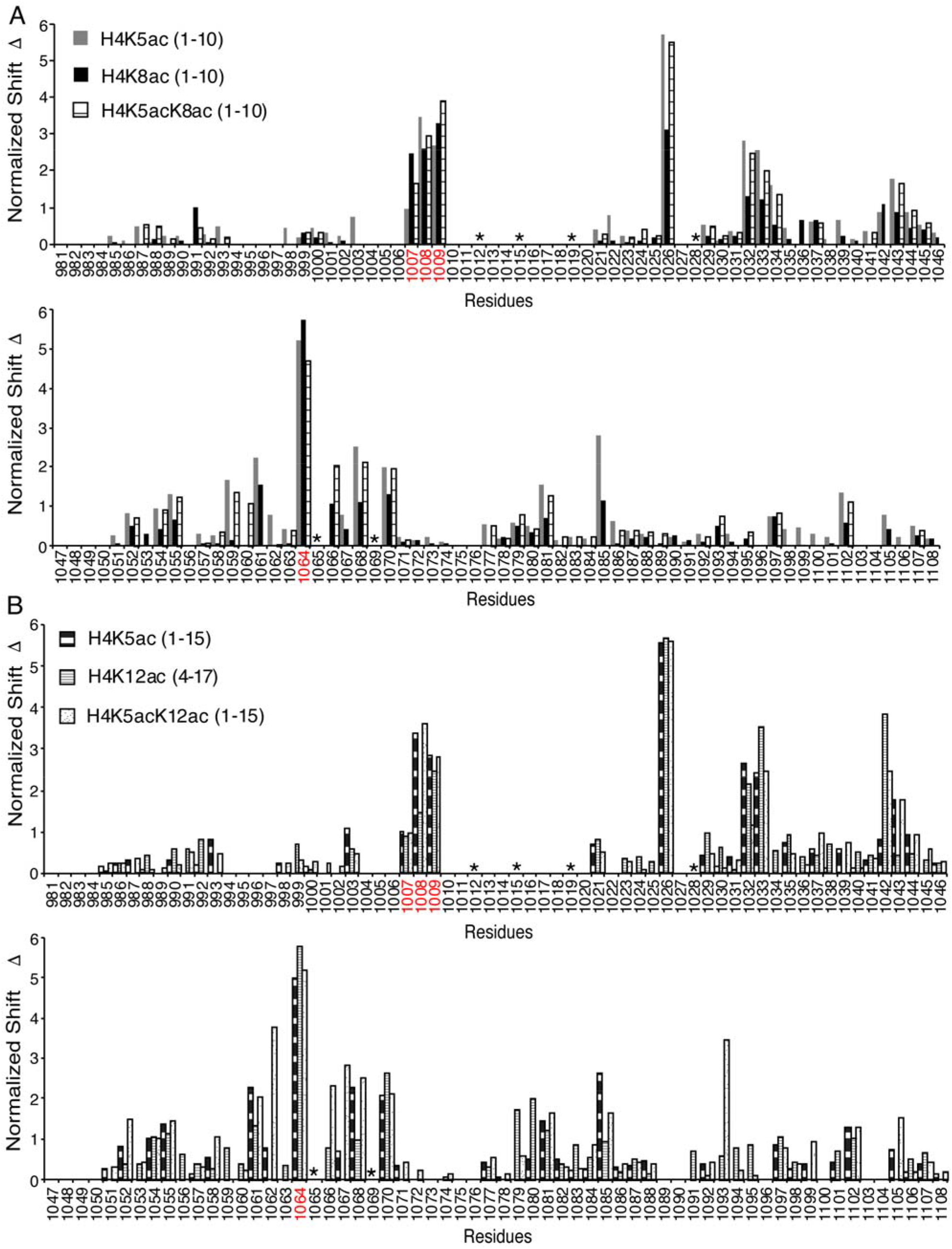
Chemical shift perturbation of ATAD2 bromodomain (BRD) residues upon interaction with histone ligands. A) Histogram showing the normalized ^1^H, ^15^N chemical shift changes in the assigned backbone amides of the ATAD2 BRD upon addition of H4K5ac (1-10), H4K8ac (1-10), and H4K5acK8ac (1-10) histone peptides. B) Histogram showing the normalized ^1^H, ^15^N chemical shift changes in the assigned backbone amides of the ATAD2 BRD upon addition of K4K5ac (1-15), H4K12ac (1-15) and H4K5acK12ac (1-15) histone peptides. Residues highlighted in red correspond to the “RVF shelf” motif and the conserved asparagine 1064. Residues highlighted by the asterisk denote prolines. The gaps in the residue axis denote unassigned amino acids.

**Figure 3:**
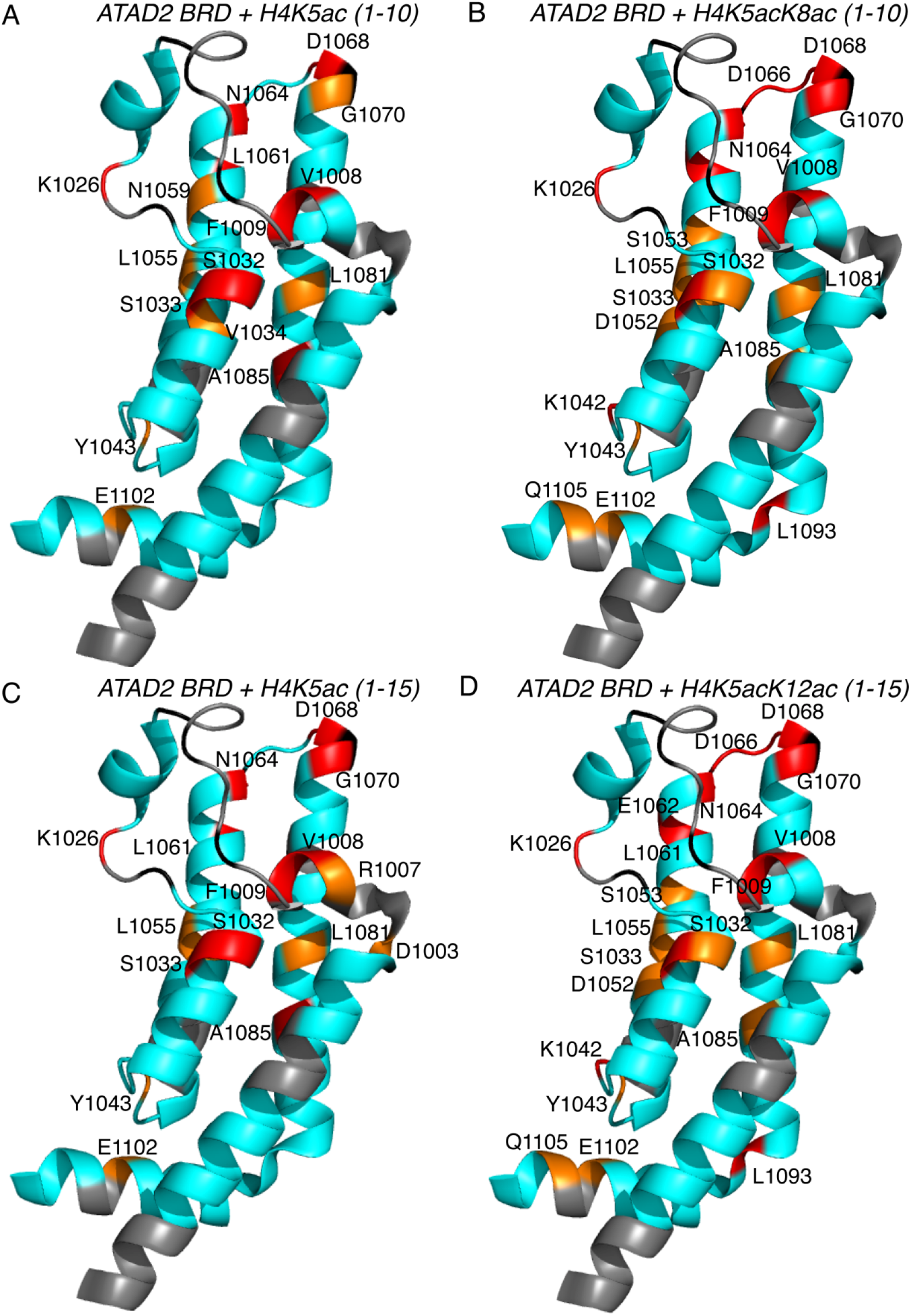
Chemical shift mapping of the ATAD2 bromodomain (BRD) highlighting the histone binding pocket. Chemical shift changes induced by addition of 1:5 molar ratio of BRD to peptides are mapped onto the crystal structure of the apo ATAD2 BRD (PDB ID: 3DAI). Residues that are not assigned are colored grey, residues exhibiting 1 or 2 standard deviations from the average chemical shift change are colored orange and red, respectively. A) Perturbations caused by addition of mono acetylated H4K5ac (1-10) histone peptide. B) Perturbations caused by addition of di-acetylated H4K5acK8ac (1-10) histone peptide. C) Perturbations caused by addition of mono-acetylated H4K5ac (1-15) histone peptide and D) Perturbations caused by addition of mono-acetylated H4K5acK12ac (1-15) histone peptide.

Comparison of the CSPs between mono- and di-acetylated histones provide new insights on the coordination of the second acetyllysine group by the ATAD2 BRD. The addition of di-acetylated histone H4K5acK8ac (1-10) resulted in more localized CSPs with stronger perturbations (>2σ) observed for residues R1007 and F1009 in the “RVF” shelf, residue A1060 in the αB helix, and residue D1066 in the BC loop of the bromodomain (**Figure 2A**). The addition of the mono-acetylated histone H4K8ac (1-10) peptide induced mostly weak CSPs (<1σ) when compared to mono-acetylated H4K5ac histone peptides). This weak binding was further confirmed by the low binding affinity observed in our ITC data (**Figure 2A**, **Table 1**). Addition of mono-acetylated histone H4K12ac (4-17) peptide, lacking the first three residues induced some CSPs with smaller perturbations for the “RVF” shelf residues and more significant CSPs for residues distributed throughout the ATAD2 BRD backbone (**Figure 2B**). Although our ITC data display similar binding affinities for the di-acetylated histone H4K5acK12ac (1-15) and the mono-acetylated histone H4K5ac (1-15) peptides, the NMR titration data displays stronger CSPs induced upon addition of the H4K5acK12ac (1-15) peptide when compared to the H4K5ac (1-15) peptide. This is especially noticeable in residues E1062, D1066 and R1067, which are located around the BC loop, and residues K1042 and L1093 that reside outside the binding pocket (**Figure 2B**, **Figure 3C, D**). It also appears that additional contacts occur between residues (D1052, S1053) lining the first acetyllysine binding pocket. Furthermore, ligand induced conformational changes in the residues lining the flexible ZA and BC loops of the bromodomain appear to assist in coordination of the second acetyllysine group. This increase in the intensity and number of perturbations induced in the presence of di-acetylated histone peptides support our hypothesis that the ATAD2 bromodomain preferentially recognizes the di-acetylated histone ligands.

### The ATAD2 bromodomain recognizes di-acetylated histone ligands as a monomer

Oligomerization is a commonly reported feature in several AAA+ domain containing proteins including ATAD2 [34]. However, there is no general agreement on the oligomeric state of the ATAD2 BRD functioning as a reader of acetyllysine. We hypothesize that ATAD2 BRD is a functionally active monomer capable of recognizing various acetyllysine marks in its monomeric state. In support of this, our ITC data display a binding stoichiometry (N: peptide-to-bromodomain ratio) of 1, indicative of a monomeric interaction between the ATAD2 BRD and various acetylated histone tail peptides (**Table 1**). This is also supported by our sedimentation velocity analysis, where the molecular mass of the ATAD2 BRD alone and in complex with acetylated histone tail peptides is approximately 15,000 Da, which further confirms that recognition of acetylated histones by the ATAD2 BRD occurs in a 1:1 stoichiometric ratio (**Figure 4**).

**Figure 4:**
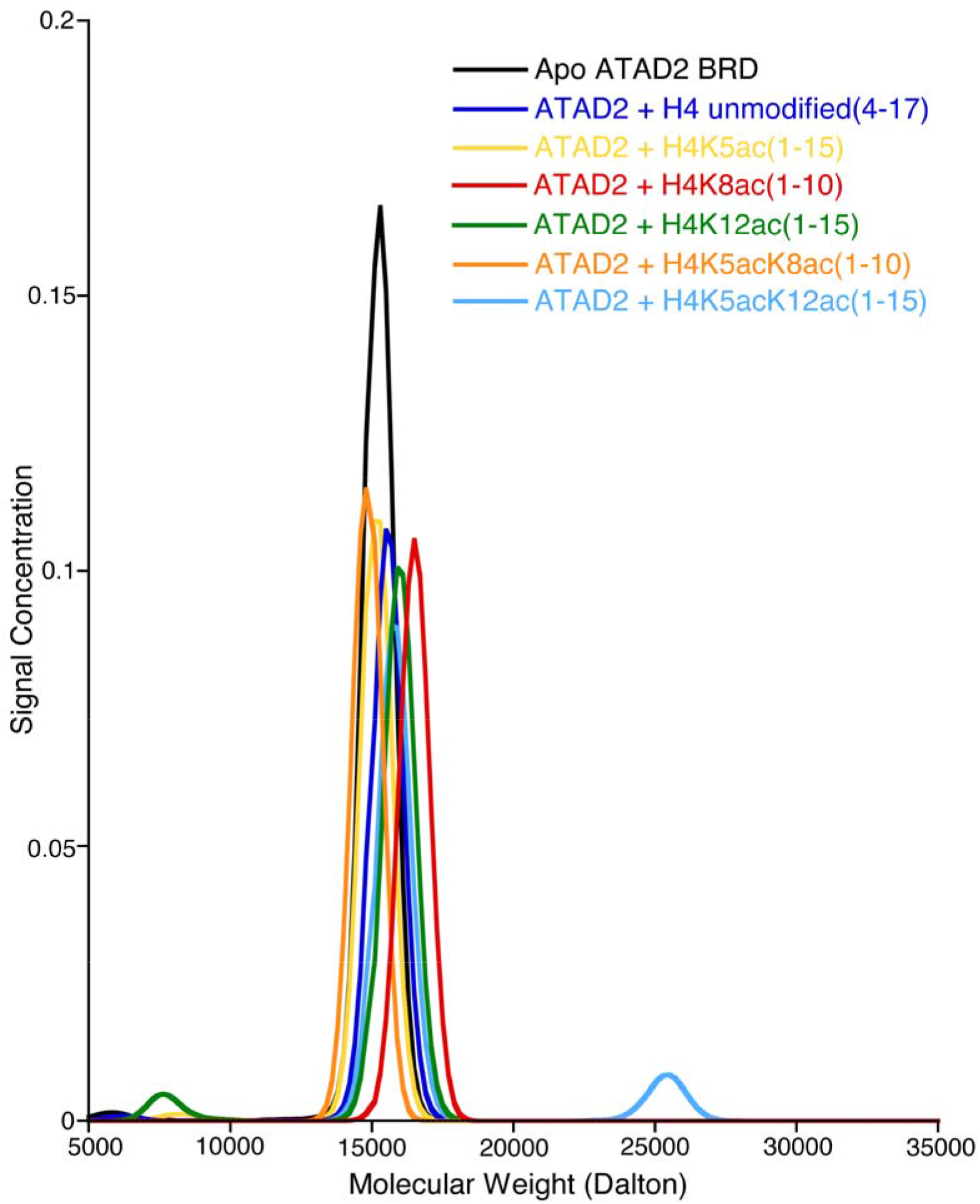
Molecular weight distribution of apo ATAD2 bromodomain (BRD) and in complex with histone H4 peptides. The molecular weights of the ATAD2 bromodomain by itself (black) and in complex with a 1:1 molar ratio with H4 unmodified (4-17) (dark blue), H4K5ac (1-15) (yellow), H4K8ac (1-10) (red), H4K12ac (1-15) (green), H4K5acK8ac (1-10) (orange) and H4K5acK12ac (1-15) (light blue) histone peptides. All molar masses are consistent with a 1:1 ratio of ATAD2 with one peptide molecule.

The oligomeric state of a protein in solution can also be determined from its rotational correlation time (τ_c_) obtained by NMR relaxation experiments [35,36]. **Table 2** gives the rotational correlation time (τ_c_) obtained from the ^15^N *T*_1_ and *T*_2_ relaxation rates of the ATAD2 BRD alone and in complex with the histone H4K5acK12ac peptide. The average *T*_1_ and *T*_2_ values of 897 ± 63 ms and 54 ± 6 ms, respectively, were obtained from the ATAD2 BRD in solution. These average *T*_1_ and *T*_2_ values were then used to estimate an overall correlation time for the protein as 12.7 ns. For a spherical, globular protein, the τ_c_ is related to the hydrodynamic radius and hence indicates its oligomeric state [37]. Thus, the slight difference in our calculated and expected τ_c_ values for the apo ATAD2 BRD can be attributed to a non-globular macromolecular shape with flexible regions, which slow down its rotational motion in solution. However, even in the presence of the di-acetylated H4K5acK12ac (1-15) histone peptides (mol. wt. = 1369.5 Da), the calculated τ_c_ of the complex remains similar to the apo ATAD2 BRD, confirming a monomeric complex in solution. Taken together, our *in-vitro* ITC, ultracentrifugation, and NMR results support that the ATAD2 BRD functions as a monomer to recognize both mono- and di-acetylated histone ligands.

**Table 2:** ^15^N, *T*_1_ and *T*_2_ relaxation rate estimates for ATAD2 bromodomain (BRD) alone and in complex with histone H4K5acK12ac (1-15) peptide. The rotational correlation time (τ_c_) is calculated from *T*_1_/*T*_2_ values via the equation: 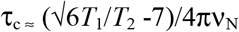

| Experimental Value | Apo ATAD2 BRD | ATAD2 BRD + H4K5acK12ac (1-15) |
| --- | --- | --- |
| $T_1$ (ms) | $897 \pm 63$ | $945 \pm 103$ |
| $T_2$ (ms) | $54 \pm 6$ | $51 \pm 13$ |
| $\tau_c$ (ns) | 12.7 | 13.4 |

### Additional residues on the N- and C-terminus of the ATAD2 BRD enhance histone tail recognition

In our efforts to improve the solubility and stability of the recombinant ATAD2 BRD, we designed a longer ATAD2 BRD construct (aa 966-1112, C1101A) containing additional N- and C-terminus residues compared to our previous construct (ATAD2 BRD aa 981-1108). Interestingly, this longer bromodomain displayed enhanced binding affinities (**Table 3**). Mono-acetylated histone H4K5ac (1-10), di-acetylated histones H4K5acK8ac (1-10), and H4K5acK12ac (1-15) exhibited almost two-fold higher binding affinities compared to the shorter ATAD2 BRD (981-1108) activity. To better understand how this longer BRD recognizes di-acetylated ligands, we grew crystals of the ATAD2 BRD (966-1112, C1101A) in complex with the histone H4K5acK8ac (1-10) peptide. The structure was solved by molecular replacement using the ATAD2 BRD structure (PDB ID: 4TT2) with the ligand removed as the search model [3]. Unfortunately, there was no electron density for the acetyllysine at position 8 in the final deposited structure (PDB ID: 7M98). However, with a resolution of 1.59 Å (see **Table 4** for the refinement statistics), we observed well-resolved electron density surrounding the K5ac group, and especially for residues Ser1, Gly2, Gly4, and acetylated Lys5 at the N-terminus of the histone H4 tail (**Figure 5C**). Coordination of the histone ligand occurs through hydrogen bonding and hydrophobic interactions (**Figure 5A, B**). Five polar contacts between the ligand and residues in the ATAD2 bromodomain are present in our structure. Y1063 is responsible for two of these contacts (**Figure 5B**). It forms hydrogen bonds using its backbone carbonyl oxygen to the backbone amide of Gly2, and through its side chain hydroxyl group to the backbone amide of Gly4. D1020 coordinates a hydrogen bond between its side chain carboxyl group and the backbone amide of Arg3 is also coordinated through a hydrogen bond between its backbone carbonyl oxygen to the side chain NH1 of R1067. As seen in other BRD structures, the conserved asparagine (N1064) coordinates the acetylation modification on residue K5ac. The carbonyl oxygen of the acetyl group is coordinated through hydrogen bonding with the side chain amide of N1064. It is also coordinated through an ordered water molecule at the base of the binding pocket that also interacts with Y1021. In addition to the polar contacts present in our structure, we have identified multiple hydrophobic contacts that add to ligand stabilization. **Figure 5A, B** shows that E1062 and P1065 line the binding pocket and interact with the N-terminus of the ligand to assist in coordinating Ser1. P1019 is found near the top of the binding pocket and can stabilize the side chain of Arg3. Members of the ZA loop, V1013, V1018, and the gatekeeper residue I1074 coordinate the acetyllysine residue, potentially by interacting with the carbon atoms in the side chain of K5ac in the binding pocket. Overall, this structure provides additional, high-resolution information for the coordination of the N-terminal histone tail by the ATAD2 bromodomain and is consistent with our biochemical and biophysical data that demonstrate recognition of di-acetylated histone ligands occurs through a monomeric binding mode.

**Table 3:** Binding affinities and stoichiometry of binding (N) obtained via ITC for acetylated histone peptide binding to the ATAD2 bromodomain (residues 966-1112).

| Histone peptide | Sequence | K<sub>D</sub> from ITC (μM) | N-value |
| --- | --- | --- | --- |
| H4K5ac (res 1-10) | SGRGKacGGKGL | 10.1 ± 1.3 | 1.021 |
| H4K5acK8ac (res 1-10) | SGRGKacGGKacGL | 15.9 ± 2.5 | 0.974 |
| H4K5acK12ac (res 1-15) | SGRGKacGGKGLGKacGGA | 15.0 ± 1.3 | 0.977 |

**Table 4:** Data collection and refinement statistics for the crystal structure of the ATAD2 bromodomain (BRD) bound to histone H4K5acK8ac (1-10).

| ATAD2 BRD in complex with histone H4K5acK8ac (1-10) |  |
| --- | --- |
| Wavelength | 1 |
| Resolution range | 44.54 - 1.59 (1.647 - 1.59) |
| Space group | P 43 2 2 |
| Unit cell | 53.56 53.56 160.35 90 90 90 |
| Total reflections | 64621 (6179) |
| Unique reflections | 32345 (3090) |
| Multiplicity | 2.0 (2.0) |
| Completeness (%) | 99.65 (97.14) |
| Mean I/sigma(I) | 18.87 (0.75) |
| Wilson B-factor | 28.00 |
| R-merge | 0.01768 (0.6612) |
| R-meas | 0.025 (0.935) |
| R-pim | 0.01768 (0.6612) |
| CC1/2 | 1 (0.495) |
| CC* | 1 (0.814) |
| Reflections used in refinement | 32340 (3090) |
| Reflections used for R-free | 1641 (139) |
| R-work | 0.1944 (0.3336) |
| R-free | 0.2195 (0.3545) |
| CC(work) | 0.967 (0.634) |
| CC(free) | 0.941 (0.595) |
| Number of non-hydrogen atoms | 1406 |
| macromolecules | 1203 |
| solvent | 203 |
| Protein residues | 140 |
| RMS(bonds) | 0.014 |
| RMS(angles) | 1.25 |
| Ramachandran favored (%) | 98.50 |
| Ramachandran allowed (%) | 1.50 |
| Ramachandran outliers (%) | 0.00 |
| Rotamer outliers (%) | 1.49 |
| Clashscore | 2.48 |
| Average B-factor | 39.75 |
| macromolecules | 38.62 |
| solvent | 46.43 |
| Statistics for the highest-resolution shell are shown in parentheses. |  |

**Figure 5:**
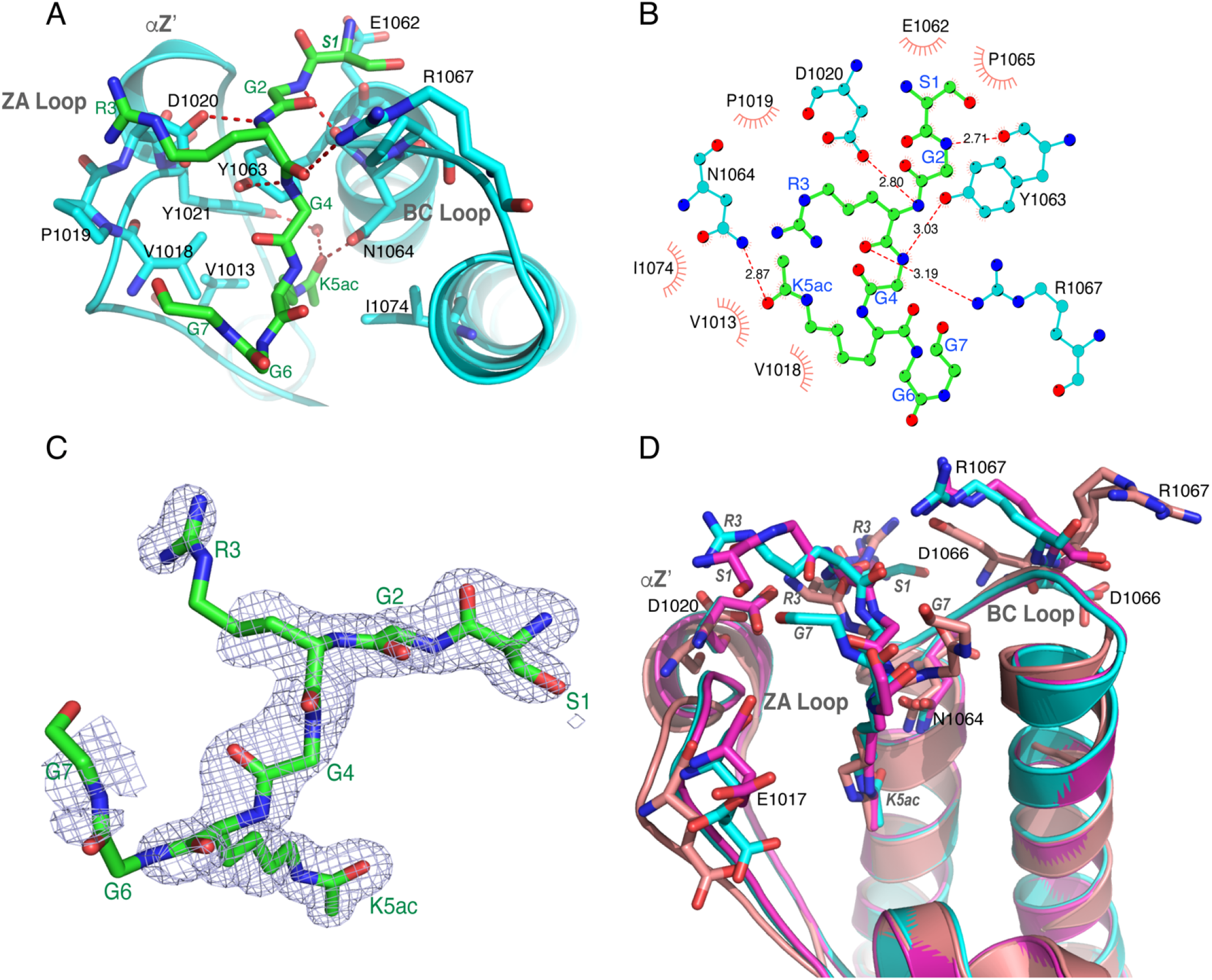
Coordination of the H4K5acK8ac (1-10) histone ligand by the ATAD2 bromodomain (BRD). A) Ligand coordination for ATAD2 BRD (cyan) with H4K5acK8ac (1-10) (green) (PDBID: 7M98). Hydrogen bonds are indicated as dashed red lines. B) LigPlot^+^ was used to depict coordination of the H4K5ac (1-10) ligand (green) by the ATAD2 BRD. Five polar contacts to D1020, Y1063, N1064, and R1067 are displayed as well as the hydrophobic interactions assisting in ligand coordination. C) Electron density for the simulated annealing composite omit map (blue) contoured at 1 σ for the histone H4K5acK8ac (1-10) ligand (green). Clear electron density is observed for the first five amino acids including the K5ac sidechain. D) Alignment of the ATAD2 BRD structures 4TT2 (magenta), 4QUU (salmon), and 7M98 (cyan). Highlighted residues are labeled in the protein backbone. The first three ligand residues display a different coordination between the overlaid structures and are labeled. Figures were generated with the PyMOL Molecular Graphics System, version 2.3, Schrödinger, LLC.

## Discussion

In this study, the ITC, AUC, and NMR relaxation studies emphasize that the ATAD2 BRD is functionally active as a monomeric chromatin reader domain that recognizes mono- and di-acetylated histones. Previous studies showed that the ATAD2 BRD is able to bind histone H4K5ac and H4K12ac ligands [3,7]. ATAD2 recognition of the H4K5acK12ac modification has been linked to its interaction with newly synthesized histones immediately following DNA replication [26], and our results indicate this di-acetylated ligand is preferred by the ATAD2 BRD over other post-translationally modified histone ligands.

Although X-ray crystal structures of ATAD2 BRD in complex with mono-acetylated histone peptides (PDB ID 4QUU, 4QUT, 4TT2) have been published previously [3,7], there is no structure for the di-acetyllysine histone in complex with ATAD2. In this study, routine 2D ^15^N-^1^H HSQC NMR experiments were used to guide the characterization of the acetyllysine binding pocket while monitoring the labeled ATAD2 BRD in solution. Backbone amide resonances of residues in the NMR spectra are sensitive to the local environment of each residue, and their positions are affected by interactions with ligands (and conformational changes), which can cause CSPs and provide useful information on the binding surface on the protein [38–41]. With majority of resonances assigned, NMR titration of mono- and di-acetylated histones into the ATAD2 BRD was valuable in defining a cluster of affected residues lining the binding pocket that are involved in the BRD-histone interaction. However, assignment of the ZA loop was difficult and limited due to high disorder in the “solvent-exposed” residues, resulting in signal broadening beyond detection [33].

Our results confirm that acetylation at K5 drives ligand binding and shows that the first three residues on the histone tail (SGR) are crucial for the recognition of the histone tail when acetylation occurs at K5 or K8. The crystal structure of the ATAD2 BRD in complex with histone H4K5ac further demonstrated that these residues make important contacts with the ATAD2 BRD residue, stabilizing acetyllysine insertion into the binding pocket. The H4K8ac (1-10) histone peptide binds with weakest affinity, similar to what was observed for other sub-family IV bromodomains, namely ATAD2B and BRPF1 [27] [42]. While the NMR CSPs confirm that the residues involved in the interaction of K5ac and K8ac are similar in ATAD2 BRD, it is the residues adjacent to the acetyllysine that control the histone peptide readout by the bromodomain and contribute to the resulting complex stability [3,33].

Histone H2A is known to have a vital role in the DNA damage response [43], with K5 acetylation playing an essential role in the cellular response to chromatin damage [44,45]. Here, we report *in-vitro* binding of the ATAD2 BRD to H2AK5ac (1-12) with an affinity comparable to the H4K5ac histone peptide (**Table 1**). Considering that the amino acid residues surrounding K5ac in the histone backbone are the same for H2AK5ac and H4K5ac, it is safe to assume that they engage similar residues in the ATAD2 BRD interaction. While the binding affinities for the preferred mono- and di-acetylated histone ligands are comparable, chemical shift mapping of di-acetylated histone binding to the ATAD2 BRD in solution, provided more dynamic and residue-specific information on factors driving di-acetylated histone tail recognition. Firstly, the smaller, more widespread CSPs (<1σ) observed for the mono-acetylated H4K5ac (1-10) ligand, were replaced by more localized and significant CSPs (>2σ) for the H4K5acK8ac (1-10) ligand. Secondly, increased perturbations were observed for residues lining the RVF shelf and BC loop in the presence of the di-acetylated H4K5acK8ac (1-10) histone compared to the H4K5ac (1-10) or H4K8ac (1-10) histone peptides. Finally, since H4K5ac (1-10), H4K8ac (1-10), and H4K5acK8ac (1-10) peptides differ only in the position of the acetylation, we propose that the increased perturbations in the presence of the di-acetylated ligand could be arising from coordination of the second acetyllysine group within the ATAD2 BRD binding pocket. This more specific interaction with the di-acetyllysine histone ligand has not been reported previously in either crystallographic or MD simulation studies. It is interesting that the two di-acetyllysine histones with same “first acetylation” (K5ac), but different “second acetylation” (K8ac/K12ac) position, result in a dissimilar CSP pattern (**Figures 2** **and** **3**), which suggests a different mode of binding for the two histones. This difference is likely the result of increased electrostatic interactions between charged ZA/BC loop residues in ATAD2 BRD with charged R/K residues on the longer histone H4K5acK12ac (1-15) tail. Structural flexibility of the histone tail peptide and high disorder in ZA and BC loops in the bromodomain further facilitates the di-acetyllysine histone recognition by providing a wider binding pocket [3,33]. Previous crystal structures have reported hydrophobic interactions mediated by side chains of gatekeeper residue I1074 and the aromatic ring of Y1063 in stabilizing the K5ac coordination [3].

We did not observe any binding of the ATAD2 BRD with the H3K14ac ligand, despite evidence in a previous crystal structure (PDB ID: 4TT4). This structure shows the ligand binding to the C-terminal region of the protein, outside of the bromodomain binding pocket, contacting R1005 and H1076. More recent screens agree with our results and show no binding of the ATAD2 BRD to the H3K14ac ligand [27], thus the structure may have been a crystal packing artifact.

Many biological processes, including ligand interaction, rely on conformational rearrangement of protein macromolecules, sometimes occurring at long-range distances from the ligand binding site [46]. In NMR, the chemical exchange of backbone-amide ^15^N nuclei are relevant in studying binding interactions, including those involving allosteric changes, with resulting chemical shifts observed as a population-weighted average of the apo and bound states [47,48]. Residue K1026 was strongly affected by addition of either mono- or di-acetylated histones (**Figures 2** **and** **3**) and while previous crystal structures of ATAD2 BRD with H4K5ac peptide (PDB ID: 4QUU and 4TT2) show no contact of this residue with the histone tail, we speculate that K1026 is an example of an allosteric conformational change during complex formation. Similarly, residue A1085 in the αC helix, is strongly affected (CSP >2σ) by addition of the mono-acetylated histone H4K5ac peptide (**Figure 3A, C**), but not by the histone H4K5acK8ac (1-10) ligand (**Figure 3B**). Since previous crystal structures do not show any interaction of this residue with the histone ligand, we suspect it is an example of an allosteric conformational change driving mono-acetylated histone binding. Likewise, residue L1093, residing at the “hinge” at the bottom of the αC helix, is strongly perturbed (CSP >2σ) when the longer di-acetylated H4K5acK12ac (1-15) peptide binds to ATAD2 BRD (**Figure 3D**). Such allosteric conformational changes have not been discussed earlier and could provide more meaningful insights into histone tail recognition. Whether they impart specificity to ATAD2 BRD coordination of acetyllysine histones is yet to be determined, and additional experiments would be needed to identify if these alternate sites could be useful for therapeutic targeting.

We also solved the crystal structure for the ATAD2 bromodomain bound to the histone H4K5acK8ac ligand at a resolution of 1.59 Å (PDB ID: 7M98). While there was missing electron density beyond ligand residue 7, we observed excellent density for residues Ser1-Lys5. We found notable differences between our high-resolution crystal structure and the previously solved ATAD2 BRD structures in complex with mono-acetylated H4K5ac ligands (PDB IDs: 4TT2 and 4QUU), solved at 2.5 Å and 1.8 Å resolution, respectively [3,7]. Differences include the orientation of residues lining the ATAD2 BRD binding pocket and the positioning of the first three ligand residues (**Figure 5D**). The loop regions of the bromodomain are the most flexible parts of its structure [33]. Therefore, it is not surprising that conformational changes differences in the position of various residues on the loop are observed for different histone ligands, especially when interacting with different acetylation marks [3,7]. In the αZ’ region of the ZA loop, residue D1020 makes a hydrogen bond to the backbone amide of ligand Arg3. This is a notable difference compared to the mono-acetylated histone complex structure in 4TT2 where ligand Ser1 is located closer to the ATAD2 D1020 residue and does not make the same hydrogen bond. Although density for the K8ac residue is missing, our structure shows that the same D1020 in ATAD2 coordinates ligand residues Ser1 and Gly2 via hydrophobic contacts. D1020 is located in a segment of the ZA loop that is the most disordered (P1015-D1020) and exhibits high conformational heterogeneity, extending into solution in the apo state and engaging multiple residues on the histone peptide [33]. The difference in positioning of the di-acetylated peptide when compared to the mono-acetylated ligands in 4TT2 and 4QUU may explain why in our structure the ZA loop region is in a slightly different orientation when compared to other structures. Also, ATAD2 residue E1017, which has been reported previously to move towards the histone peptide in the bound state in order to make potential salt bridges with the K8 residue, is found in two alternate conformations in our structure with the H4K5acK8ac ligand [33]. Whereas in the structures of ATAD2 bound to mono-acetyllysine only one conformation is observed [3,7].

The BC loop is another flexible region in the bromodomain binding pocket that contains the conserved asparagine, as well as several other important contacts that were identified in our analysis. While there is no significant difference in the orientation of residue N1064 in the ATAD2 BRD:H4K5acK8ac structure compared to previous structures, the side chain position allows it to reach deeper into the binding pocket to make a closer hydrogen bond contact (2.87 Å) than in the 4TT2 (3.4 Å) and 4QUU (2.92 Å) structures. Importantly, the side chains of residues D1066 and R1067, also in the BC loop, are in slightly different positions. This allows for the side chain of R1067 to make a hydrogen bond contact with the backbone carbonyl group of Arg3 and supports our NMR data showing a CSP for D1066.

Overall, differences between our 7M98 structure and the 4TT2 and 4QUU structures appear to reflect the flexibility of the ZA and BC loops. Although they are subtle, these variations may result distinct molecular interactions required for mono- and di-acetyllysine coordination. Where the latter promotes stabilizing conformations of residues in the ATAD2 BRD binding pocket, rigidifying the histone peptide backbone. Future studies are needed to determine the structure of ATAD2 bound to a di-acetylated ligand to further illustrate the molecular details of this interaction.

### Histone H4 tail recognition by ATAD2 BRD differs from its ATAD2B BRD paralogue

Since the AAA-containing bromodomain, ATAD2B, is known to share high amino-acid sequence similarity (97%) and identity (74%) with the ATAD2 BRD [49], we expected these two BRDs to engage mono- and di-acetylated histone tail residues in a similar manner. Interestingly, although the ATAD2/B BRDs share a similar subset of PTM histone ligands, the ATAD2B BRD had a much broader binding specificity, recognizing 39 histone ligands compared to 11 for the ATAD2 BRD [27]. Considering that recognition of acetylated histones is heavily influenced by the peptide backbone sequence, it was not surprising that mono-acetylated H2A and H4 tail peptides displayed similar binding affinities [27]. The only exception was with histone H4K5ac (1-15), which binds more tightly to the ATAD2B BRD than to the ATAD2 BRD. We posit that tighter binding to the ATAD2B BRD arises from additional contacts between charged residues on the ZA helix and BC loops, which coordinate the H4K5ac (1-15) peptide [27]. Likewise, the majority of the di-acetylated histones also displayed similar binding affinities for the ATAD2/B BRDs. However, the NMR titration data in this study highlight key differences in the residues engaging the di-acetyllysine histone coordination when compared to previously published ATAD2B BRD data [27]. For example, similar residues were perturbed for the two di-acetyllysine histone ligands H4K5acK8ac (1-10) and H4K5acK12ac (1-15) in the ATAD2B BRD [27], whereas in the ATAD2 BRD more specific and unique CSP were displayed by residues involved in di-acetylated histone coordination. Also, in the ATAD2B BRD, mutation of the conserved asparagine (N1038A) reduced the binding affinity significantly, but did not abrogate binding to acetylated histones. However, in the ATAD2 BRD, the corresponding N1064 plays a more significant role in coordination of the acetyllysine group, displaying one of the highest CSPs. Furthermore, mutation of N1064A resulted in complete loss of ligand binding in the ATAD2 BRD. This is in line with previous studies showing that N1064 contacts the K5ac group through a direct hydrogen bond. Despite high sequence similarity, the residues engaging the di-acetyllysine histones vary within ATAD2/B BRDs. While the ATAD2B BRD showed a similar CSP pattern overall for coordination of either mono- or di-acetylated histones, the ATAD2 BRD displays relatively more unique and significant CSPs upon interaction with mono- versus di-acetylated histone ligands. It is interesting to note that in the ATAD2 BRD, the “RVF” shelf and BC loop residues (E1062, D1066 and R1067) were most significantly affected in the presence of di-acetylated histones, while corresponding residues in ATAD2B BRD: the NIF shelf, E1036, D1040 and K1041 (in the BC loop) displayed only weak perturbations (> 1σ) [27]. Additionally, while both ATAD2/B have a similar charge density around the binding pocket, it appears that the ATAD2 BRD engages di-acetylated histones via charged residues, while the ATAD2B BRD relies more heavily on contacts via hydrophobic residues. Residues (K1026, A1085 and L1093), which display allosteric changes when ATAD2 BRD binds to mono- or di-acetylated histones, show either weak or no CSPs upon ATAD2B BRD ligand binding. Although these BRDs share high sequence identity, a detailed analysis of their binding pockets emphasizes subtle, yet significant differences in the residues engaging acetylated histones, which could be useful in the design of more selective inhibitor molecules aiding cancer therapeutics. Overall, our study highlights subtle differences in mono-versus di-acetylated histone tail coordination by ATAD2 BRD residues, including those that undergo allosteric changes upon ligand binding.

## Experimental Procedures

### Plasmid construction

The wild type human ATAD2 BRD cDNA was a gift from Dr. Nicola Burgess-Brown (Addgene plasmid #38916). Residues 981-1108 encoding the 128 amino acid ATAD2 BRD were PCR amplified and re-cloned using the Gateway Cloning technology (Invitrogen) into the pDest15 vector (Invitrogen) with an N-terminal glutathione transferase (GST) tag followed by a PreScission Protease site (GE Healthcare). The codon optimized cDNA for the ATAD2 BRD was designed by BioBasic and subsequently amplified by PCR and cloned into the pDest15 vector. The ATAD2 BRD mutants N1064A, I1074A, and I1074Y were created using the QuikChange^®^ II XL Site-Directed Mutagenesis Kit (Stratagene) using previously described protocols [50]. DNA sequences were verified (University of Vermont Cancer Center Advanced Genome Technologies Core, Eurofins) and then transformed into *E. coli* Rosetta 2(DE3) pLysS competent cells (Novagen) for non-codon optimized sequences, and into *E. coli* BL21(DE3) pLysS chemically competent cells (Invitrogen) for codon optimized (CO) sequences.

The ATAD2 long construct (966-1112, C1101A) was cloned into the pGEX-6P-1 plasmid with an N-terminal GST tag and precision protease cleavage site after codon optimization by DAPCEL and synthesis by GenScript. The plasmid was then transformed into *E. coli* BL21(DE3) pLysS competent cells (Novagen) for protein expression.

### Protein expression and purification of the ATAD2 bromodomain

*E. coli* cells containing the GST-tagged ATAD2 WT (981-1108), ATAD2 (981-1108) with a N1064A/I1074A/I1074Y mutation, or the ATAD2 (966-1112, C1101A) plasmids, were grown in 4 L of Terrific Broth (TB) or ^15^N ammonium chloride or ^13^C-glucose supplemented minimal media at 37°C and shaking at 225 RPM. The temperature of the culture was dropped to 20 °C for 1 h upon reaching an OD_600_ of 0.8. After 1 h at 20 °C, protein expression was then induced by addition of 0.5 mM isopropyl ß-D-1-thiogalactopyranoside (IPTG), and cells were harvested by pelleting after 20 h of incubation at 20 °C and shaking at 225 RPM. The bacterial cell pellet was weighed and re-suspended in 200 mL of lysis buffer (50 mM Tris pH 7.5, 500 mM NaCl, 5% glycerol, 0.05% Nonidet P-40) with 2 mL of lysozyme and a protease inhibitor tablet (Thermo Scientific). The cells were then lysed by sonication, and the lysate was cleared by centrifugation at 10,000 RPM for 20 min. The supernatant was added to 10 mL glutathione agarose resin (Thermo Scientific) and incubated on ice at 4 °C while rotating for a minimum of 2 h. After incubation, the suspension was centrifuged for 5 min at 1,000 x g to collect the agarose beads bound to protein. Once collected, the beads were washed in wash buffer (50 mM Tris pH 7.5, 500 mM NaCl, 5% glycerol) 3 times and moved into a 25 mL Econo-Column Chromatography Column (Bio-Rad), where they were further washed with a minimum of 200 mL of wash buffer. The GST tag was cleaved by the addition of PreScission Protease (GE Healthcare) with rocking incubation overnight at 4 °C. The protein was then eluted with wash buffer supplemented with 1 mM dithiothreitol (DTT). The concentration of the ATAD2 BRD was determined either by NanoDrop (Thermo Scientific) measurement of the absorption at 280 nm with the extinction coefficient of 7450 M^−1^ cm^−1^, or by using the Pierce BCA Assay Kit (Thermo Scientific) from the absorption at 550 nm. SDS-PAGE gels stained with GelCode Blue Safe protein stain (Thermo Scientific) confirmed the purity of the ATAD2 BRD protein samples.

### Histone peptide synthesis

Histone tail ligands were synthesized at the Peptide Core Facility at the University of Colorado Denver, Biomatik Inc., and/or GenScript either as unmodified histones or designed with specific acetyllysine modifications incorporated. Each peptide contains a free N-terminus and an amidated C-terminus. The peptides were supplied as TFA salt and refined to greater than 98% purity via HPLC. Their chemical identities were confirmed by mass spectroscopy before use. Sequence information for each of the peptides used in this study is shown in **Table 1**.

### Isothermal titration calorimetry

Purified protein was concentrated to a volume of 3 mL and dialyzed into ITC buffer (150 mM NaH_2_PO_4_ and 150 mM NaCl) for 48 h. ITC experiments with histone peptides were carried out at 5 **°**C using a MicroCal iTC200 (Malvern Panalytical). The ATAD2 bromodomain and histone peptide samples were diluted with ITC buffer (150 mM NaH_2_PO_4_ and 150 mM NaCl). Calorimetric titration was performed by titrating the histone tail ligands at a concentration of 5 mM into 0.2 mM of the ATAD2 bromodomain protein in the ITC sample cell in a series of 20 injections, the first at 0.5 μL, and the rest at 2.0 μL with time intervals of 150 s. The stirring speed was set at 750 RPM. After excluding the first injection, the ITC binding isotherms were integrated using a Marquandt nonlinear least-squares analysis according to a 1:1 binding model assuming a single set of identical binding sites with the Origin 7.0 program (OrginLab Corporation). The raw data were corrected for non-specific heats of dilution determined by the magnitude of the peaks appearing after the system reaches complete saturation. The obtained heats of dilution were analyzed according to a one set of sites binding model and used to calculate binding affinities, N values, and thermodynamic values. Experiments showing no binding were repeated in duplicate, while those showing a binding interaction were completed in triplicate. The mean KD was calculated from the average of the three runs, and the standard deviation was calculated from the mean.

### Analytical ultracentrifugation

For analytical ultracentrifugation (AUC), the bromodomain was purified further using fast-protein liquid chromatography on an AKTA Purifier UPC 10 (GE Healthcare), equipped with a HiPrep 16/60 Sephacryl S-100 High Resolution column (GE Healthcare), after being equilibrated with AUC Buffer (50 mM NaH2PO4, 50 mM NaCl, and 1 mM tris(2-carboxyethyl)phosphine (TCEP). Fractions were collected using a Frac-920 fraction collector (GE Healthcare). Purified protein samples were prepared at an A_280_ of approximately 0.6 and subsequently checked using the Eppendorf Biophotometer Plus Spectrophotometer to get accurate A_280_ readings for the data calculations. Histone peptides studied included H4 unmodified (4-17), H4K5ac (1-15), H4K8ac (1-10), H4K5acK8ac (1-10), H4K5acK12ac (1-15), and H4K12ac (1-15) that were added in a 1:1 molar ratio. Samples were kept at 4 °C until the experiment. The ATAD2 bromodomain alone, and the ATAD2 bromodomain combined with unmodified histone peptides, were included as controls. All measurements were made at 20 ºC by UV intensity detection in a Beckman Optima AUC analytical ultracentrifuge at the Center for Analytical Ultracentrifugation of Macromolecular Assemblies at the University of Texas Health Science Center at San Antonio, using an An50Ti rotor and standard 2-channel epon centerpieces (Beckman-Coulter). All samples were measured in the same run, and at multiple wavelengths (278 nm, 280 nm and 282 nm).

The data was analyzed with UltraScan-III ver. 4.0, release 2480 [51]. Hydrodynamic corrections for buffer density and viscosity were estimated by UltraScan to be 1.006 g/mL and 1.0239 cP. The partial specific volume of ATAD2 (0.7331 mL/g) was estimated by UltraScan from the protein sequence as previously outlined [52]. Sedimentation velocity (SV) data were analyzed according to the method described in [53]. Optimization was performed by 2-dimensional spectrum analysis (2DSA) [54,55] with simultaneous removal of time- and radially-invariant noise contributions [56]. After inspection of the 2DSA solutions, the fits were further refined using the parametrically constrained spectrum analysis (PCSA) [57], coupled with Monte Carlo analysis to determine confidence limits for the determined parameters [58]. The calculations are computationally intensive and were carried out on high-performance computing platforms [59]. All calculations were performed on the Lonestar5 cluster at the Texas Advanced Computing Center (TACC) at the University of Texas at Austin and on XSEDE clusters at TACC and at the San Diego Supercomputing Center.

### Nuclear magnetic resonance spectroscopy

Protein for NMR experiments was expressed and purified as previously described [60]. Uniformly labeled ^15^N ATAD2 bromodomain samples were prepared at 0.5 mM in NMR buffer containing 20 mM Tris-HCl pH 6.8, 150 mM NaCl, 10 mM DTT, and 10% D_2_O for the chemical-shift perturbation experiments. Titration mixtures containing 35 μL of the protein and peptide were made at molar concentration ratios of 1:0, 1:0.25, 1:0.5, 1:1.1, 1:2.5, and 1:5 for unlabeled histone tail peptides with specific acetyllysine modifications H2AK5ac (1-12), H4K5ac (1-15), H4K5ac (1-10), H4K5ac (4-17), H4K8ac (1-10), H4K8ac (4-17), H4K12ac (4-17), H4K5acK8ac (1-10), H4K5acK12ac (1-15), (H3K14ac (8-19), H3 unmod, and H4 unmod. The 2D ^15^N-HSQC (heteronuclear single quantum interference) experiments were performed at the National Magnetic Resonance Facility at Madison (NMRFAM). All HSQC experiments were run at 25 **°**C on a 600 MHz Bruker AVANCE III spectrometer with a z-gradient 1.7 mm TCI probe and the data collected with ^1^H at 4.475 ppm and ^15^N at 116 ppm radio-frequency pulses applied. 1024 × 128 complex data points with spectral widths of 16 ppm and 30 ppm, respectively, were collected from the ^1^H and ^15^N dimensions, with 32 scans per FID and an inter-scan delay of 1.0 second. This resulted in a total experiment time of approximately 160 min for each HSQC spectrum.

The ^15^N longitudinal (*T*_1_) and the ^15^N transverse (*T*_2_) relaxation experiments measured for the ATAD2 bromodomain alone and in complex with the H4K5acK12ac (1-15) peptide. These relaxation measurements were performed by recording a series of standard two-dimensional (2D) ^1^H, ^15^N spectra that include either an inversion recovery relaxation delay for *T*_1_, or a Carr-Purcell-Meiboom-Gill (CPMG) relaxation period for *T*_2_ [61]. Multiple 2D spectra were acquired in an interleaved manner using different relaxation delays that spanned from 40 ms to 1.6 sec for *T*_1_, and from 10 ms to 160 ms for *T*_2_. All spectra were recorded on a 600 MHz Varian VNMRS spectrometer equipped with a z-gradient 5.0 mm cryogenic probe at the National Magnetic Resonance Facility at Madison (NMRFAM). The temperature of the sample was regulated at 25°C throughout the experiments. For each 2D spectrum 1024×128 complex points were recorded for the ^1^H and ^15^N dimensions, respectively. Each FID was accumulated with 16 transients and a recovery delay of 1.5 s was used in between FIDs. The spectra were processed with NMRPipe and analyzed using NMRFAM-Sparky [62,63]. The relaxation time constants for each backbone amide were calculated in NMRFAM-Sparky by fitting the decay in the intensity of the corresponding peaks to an exponential function. The overall rotational correlation time (τC) for the protein was estimated from the ratio of *T*_1_ and *T*_2_ ^15^N relaxation times, and the nuclear frequency (νN) [61].

NMR assignments ATAD2 bromodomain were generously provided by Bayer AG and transferred onto our apo-ATAD2 bromodomain (981-1108) spectra using SPARKY [63,64] (**Supplementary Figure S3**). Due to protein instability and precipitation during NMR data acquisition, the current NMR assignment stands at 75% complete (95/127 residues). CSPs from the final titration point were calculated and normalized as described by Williamson [40], using the equation:

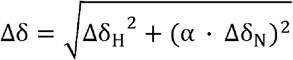

Based on our data, our scaling factor for most residues was calculated to be 0.146, and 0.2 was used for glycine. The CSPs were plotted as a bar graph to identify residues that moved significantly in response to the addition of specific histone ligands. These were calculated using the standard deviation (Δ ppm/ Standard deviation ppm, where stdev = 0.0981 ppm) of the Euclidean chemical change of the residues as described previously [27].

### Circular dichroism spectroscopy

Circular Dichroism (CD) spectroscopy was used to confirm the secondary structure of wild-type ATAD2 bromodomain and mutants N1064A, I1074A, and I1074Y. These spectra were recorded on a JASCO J-815 CD Spectrometer (JASCO, Japan) at 25 °C using a 1.6 cm cell with a 1 mm path-length (Starna Cells, Inc.). ATAD2 bromodomain wild type (WT) or mutant proteins were dialyzed in 50 mM NaH_2_PO_4_–Na_2_HPO_4_ pH 7.0, 50 mM NaCl and diluted to 20 μM. CD spectra were measured from 195 to 260 nm. Two spectra were measured at a scan rate of 20 nm/min with a bandwidth of 1 nm and averaged for each protein sample. Processed data were baseline corrected against a spectrum of the buffer alone. Spectra were analyzed using the K2D3 structure prediction software to determine the percent α-helical and β-sheet content [65].

### X-ray Crystallography

For crystallization trials the ATAD2 BRD was further purified by gel-filtration using a HiPrep 16/60 Sephacryl S-100 High Resolution column (GE Healthcare). The purified ATAD2 BRD 966-1112, C1101A protein was incubated with the histone H4K5acK8ac (1-10) peptide on ice at a 1:10 molar protein: peptide ratio. Crystallization screens were set up in 96-well, sitting drop VDX plates (Hampton Research), with each drop (2 μL) consisting of 1 μL of the protein-peptide complex plus 1 μL of mother liquor solution, with a 25 μL total reservoir volume. Initial crystals of the ATAD2 BRD in complex with the histone H4K5acK8ac (1-10) peptide grew from the Hampton Research Index screen in conditions 17-18 (1.26 M Sodium phosphate monobasic monohydrate, 0.14 M potassium phosphate dibasic, pH 5.6 and 0.49 M Sodium phosphate monobasic monohydrate, 0.91 M Potassium phosphate dibasic, pH 6.9) at 4 °C. The crystals were then reproduced in hanging drops with 24-well VDX plates (Hampton Research) by mixing 1 μL of the protein-peptide solution with 1 μL of mother liquor in each 2 μL drop over a 500 μL volume of reservoir solution. After screening against pH and salt concentrations, the optimal crystallization condition for the complex was found to be 2.0 M sodium phosphate monobasic monohydrate and 1.99 M Potassium phosphate dibasic, pH 6.6.

The crystals were harvested and mounted using B1A (ALS style) reusable goniometer bases inserted with a 300 mM Dual Thickness Microloop LD built on 18 mm / SPINE length rods (pins) from MiTeGen. The crystals were cryoprotected by adding 40% glycerol in mother liquor to a final concentration of 24% glycerol in the drop, and then flash freezing in liquid nitrogen. The data for the crystal structure was collected at the Advanced Light Source at the Lawrence Berkeley National Lab on 4.2.2 Beamline, equipped with an RDI CMOS-8M detector. The diffraction data was processed using XDS [66], and the structure was solved by molecular replacement using PHASER [67]. The molecular replacement staring model was derived from the ATAD2 BRD-H4K5ac (1-20) structure (PDB ID: 4TT2), with the histone ligand removed [3]. Once a solution was found, a structural model of ATAD2B in complex with the histone H4K5acK8ac ligand was built in COOT [68], followed by iterative rounds of refinement and rebuilding in PHENIX and COOT [69]. The final structure at 1.59 Å resolution was deposited into the Protein Data Bank under PDB ID: 7M98 after validation using MolProbity [70].

## Supporting information

Evans CM 2021 Supplemental Information

## Accession Codes

The crystal structure of the ATAD2 bromodomain in complex with histone H4K5acK8ac (1-10) has been deposited in the Protein Data Bank (PDB ID: 7M98). The authors will release the atomic coordinates upon article publication.

## Acknowledgements

Funding for this research was provided by a National Institutes of Health grants, NIGMS R01GM129338, NIGMS R15GM104865, and NCI P01CA240685 to KCG. This research was also supported by the University of Vermont Cancer Center. CME was the recipient of an ACPHS Student Summer Research Award in 2016, and an ACPHS graduate research assistantship from 2017-2018. Analytical ultracentrifugation experiments were performed at the Center for Analytical Ultracentrifugation of Macromolecular Assemblies (CAUMA) at UT Health San Antonio, which is supported by NIH grant 1R01GM120600 (BD). UltraScan supercomputer calculations were supported through an NSF/XSEDE allocation grant TG-MCB070039N (BD). Equipment was purchased with funds from the University of Texas grant TG457201 (BD). This study made use of the National Magnetic Resonance Facility at Madison, which is supported by NIH grant P41GM136463, old number P41GM103399 (NIGMS) and P41RR002301. Equipment was purchased with funds from the University of Wisconsin-Madison, the NIH P41GM103399, S10RR02781, S10RR08438, S10RR023438, S10RR025062, S10RR029220), the NSF (DMB-8415048, OIA-9977486, BIR-9214394), and the USDA.

## Author Contributions

Conceptualization, K.C.G., C.M.E., M.P., and B.D.; validation, C.M.E, M.P., K.L.M. and K.C.G.; investigation, C.M.E., K.L.M., M.T., G.C., C.C., S.H., M.G., L.W., S.C., J.C.G., J.C.N.; writing—original draft preparation, C.M.E.; writing—review and editing, C.M.E., M.P., K.L.M., G.C., S.H., M.G., B.D., J.L.M, and K.C.G.; supervision, K.C.G.; funding acquisition, B.D., J.L.M., and K.C.G. All authors have read and agreed to the published version of the manuscript.

