## Supplementary material for "Coordination of di-acetylated histone ligands by the ATAD2 bromodomain": Evans CM 2021 Supplemental Information

**Supplementary Table S1:** Dissociation constants and N-values of the interactions of the ATAD2 mutant bromodomains with a selection of histone tail peptides.

| **ATAD2 Bromodomain Mutant** | **Ligand** | **K_D_ (μM)** | **N-value** |
| --- | --- | --- | --- |
| Wild Type | H4 unmod | No binding | - |
|  | H4K5ac (1-15) | **39.4 ± 6.0** | 1.040 |
|  | H4K12ac (4-17) | 95.1 ± 13.1 | 1.170 |
|  | H4K5acK8ac (1-10) | 33.2 ± 5.9 | 0.938 |
|  | H4K5acK12ac (1-15) | 28.4 ± 1.3 | 0.910 |
| N1064A | H4 unmod | No binding | - |
|  | H4K5ac (1-15) | No binding | - |
|  | H4K12ac (4-17) | No binding | - |
|  | H4K5acK8ac (1-10) | No binding | - |
|  | H4K5acK12ac (1-15) | No binding | - |
| I1074A | H4 unmod | No binding | - |
|  | H4K5ac (1-15) | No binding | - |
|  | H4K12ac (4-17) | No binding | - |
|  | H4K5acK8ac (1-10) | No binding | - |
|  | H4K5acK12ac (1-15) | No binding | - |
| I1074Y | H4 unmod | No binding | - |
|  | H4K5ac (1-15) | No binding | - |
|  | H4K12ac (4-17) | No binding | - |
|  | H4K5acK8ac (1-10) | No binding | - |
|  | H4K5acK12ac (1-15) | No binding | - |

**Supplementary Table S2:** Secondary structure of the mutant ATAD2 bromodomain (BRD) proteins as measured by circular dichroism and calculated by K2D3.

| **ATAD2 BRD Mutant** | **% α-helix** | **% β-strand** |
| --- | --- | --- |
| **Wild Type** | **92.56%** | **0.47%** |
| **N1064A** | **92.56%** | **0.49%** |
| **I1074A** | **92.56%** | **0.49%** |
| **I1074Y** | **91.82%** | **0.50%** |
| **R1005A** | **91.96%** | **0.50%** |
| **H1076A** | **91.96%** | **0.50%** |
| **R1077A** | **91.82%** | **0.50%** |
| 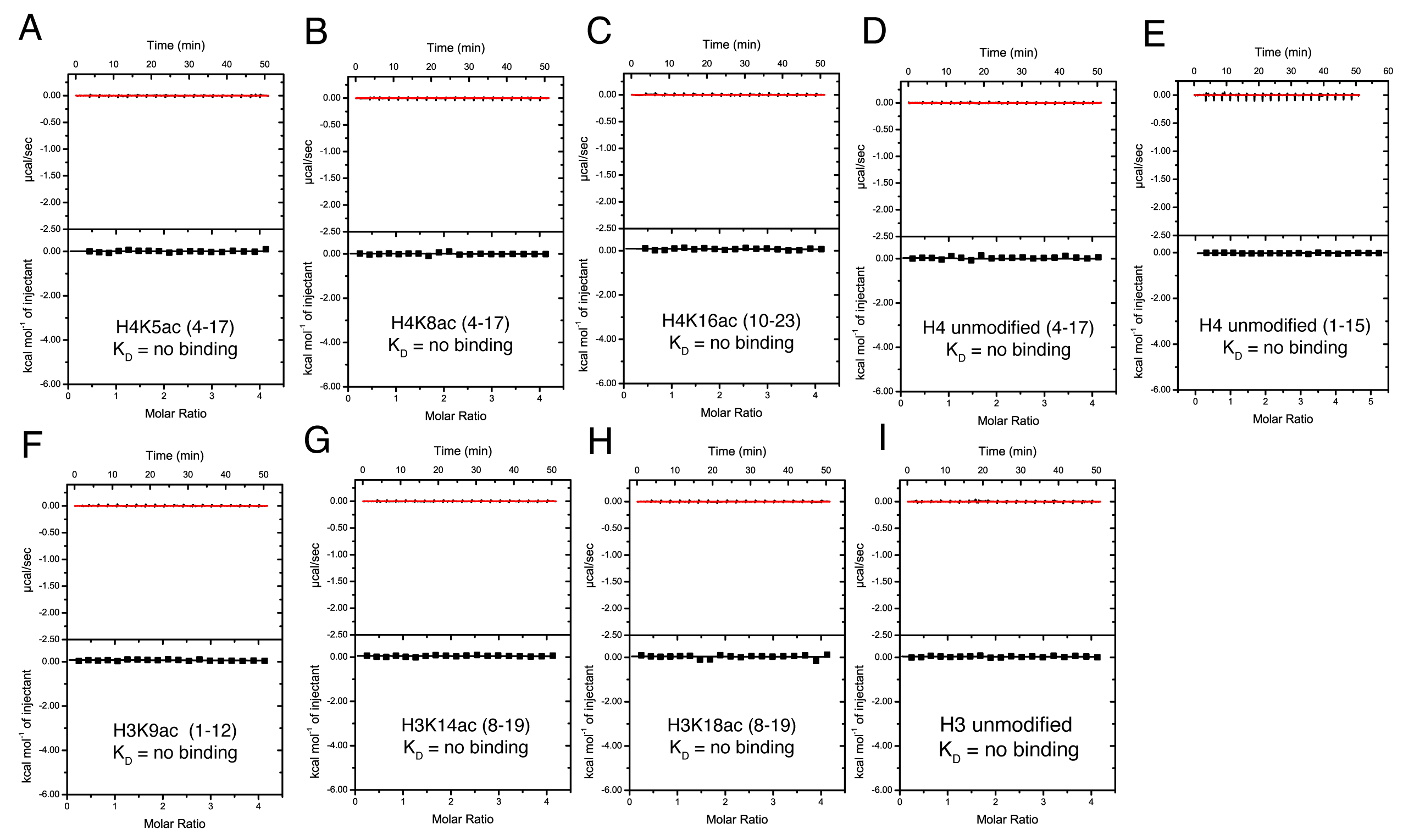 | | |

**Supplementary Figure S1:** Exothermic isothermal titration calorimetry (ITC) enthalpy plots for the binding of the ATAD2 bromodomain with histone peptides. A-I) Exothermic ITC enthalpy plots for binding of ATAD2 bromodomain with histone peptides. Calculated binding constants and peptides are indicated for each trace.

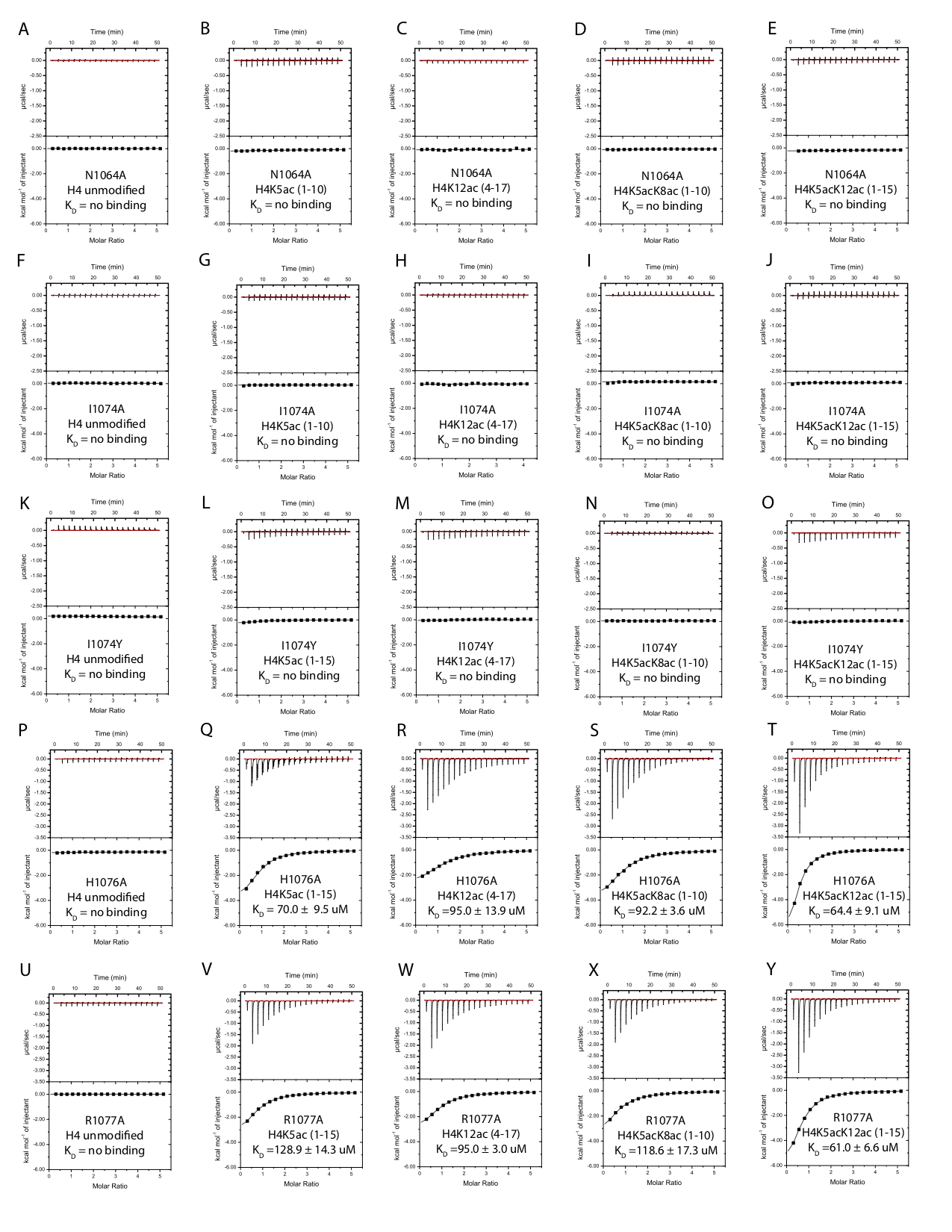

**Supplementary Figure S2:** Exothermic isothermal titration calorimetry (ITC) enthalpy plots for the binding of the ATAD2 bromodomain mutants with histone peptides. Histone ligands tested include H4 unmodified (4-17), H4K5ac (1-15), H4K12ac (4-17), H4K5acK8ac (1-10), and H4K5acK12ac (1-15).

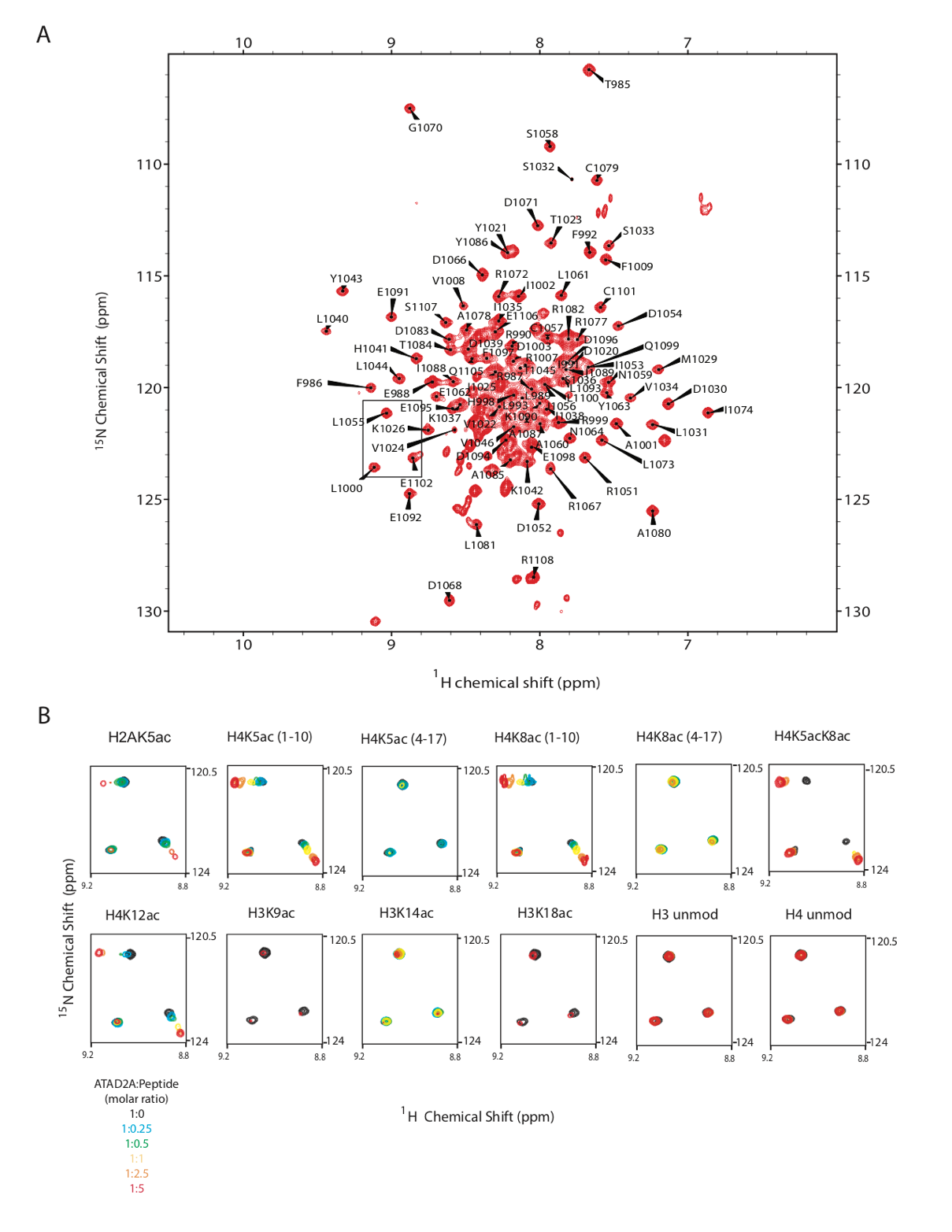

**Supplementary Figure S3:** Nuclear magnetic resonance assignment and HSQC titration data for the ATAD2 bromodomain. A) 2D ^1^H-^15^N HSQC spectrum of the ^15^N-labelled ATAD2 bromodomain showing backbone-amide assignments B) Triad of residues in the inset A contain superimposed ^1^H-^15^N HSQC spectra of the ATAD2 BRD acquired with increasing concentrations of the acetylated histone tail peptides, as indicated. The spectra are color-coded according to the ratio of protein to peptide as given in the lower left corner.
